# Diversity of cave Phlebotomines (Diptera: Psychodidae) from a Colombian cave

**DOI:** 10.1101/2022.01.07.475242

**Authors:** Manuela Velazquez, Adam M. M. Stuckert, Rafael J. Vivero, Daniel R. Matute

## Abstract

Sandflies are vector species of *Leishmania*, among many other pathogens, with a global distribution and a variety of ecological niches. Previous samplings have found that karstic formations (i.e., caves and folds formed by the erosion of limestone) serve as a natural habitat to sandfly species. The majority of samplings of cave sandfly diversity have occurred in Brazil and to date none have studied the species composition in a cave in the Northern Andes. We collected sandflies in the Cave “Los Guácharos”, in the state of Antioquia, Colombia. The sampling was carried out during two consecutive nights in September 2019. CDC-type light traps were installed inside the cavern and in other surrounding karst systems (caves and folds). In total, we identified 18 species of sandfly from the cave and surrounding karst systems, including three new records for Colombia (*Bichromomyia olmeca nociva, Brumptomyia brumpti*, and *Warileya leponti*), and provide the first karstic reports for four other species (*Lutzomyia gomezi, Lutzomyia hartmanni, Pintomyia ovallesi*, and *Psychodopygus panamensis*). We then used the results of our survey and published literature to test two hypotheses. First, that sandfly diversity in Neotropical caves is richest nearer to the equator and second that there is a phylogenetic signal of karstic habitat use in sandflies. Counter to our predictions, we found no evidence that diversity follows a latitudinal gradient. Further, we find no evidence of a phylogenetic signal of karstic habitat use, instead finding that the use of caves likely evolved multiple times across several genera. Our results highlight the importance of a wide sampling to understand the natural habitat of sandflies and other disease vectors.

## INTRODUCTION

The family Psychodidae is a speciose group of dipterans that is globally distributed. The family encompassess seven different subfamilies and has over 2,500 species (Zhang 2011; Bejarano and Estrada 2016). The majority of the species diversity is encompassed by two subfamilies. The subfamily Psychodinae (Wagner 2004; Espíndola *et al*. 2012) commonly referred to as moth flies or drain flies, which includes 4,000 species, many of which are human commensals (Cordeiro and Wagner 2018; Munstermann 2019). The second subfamily, Phlebotominae, includes 900 species, of which over 90 are vectors of diseases of humans and other animals alike (Killick-Kendrick 1990; Ready 2013). These species are commonly referred to as sandflies and are the main vectors of Leishmaniasis in the tropics and subtropics. Just for Leishmaniasis, one of the diseases transmitted by members of the subfamily Phlebotominae, more than 12 million people are infected and over 2 million new cases are reported annually. The number of recorded deaths due to *Leishmania* infection per year is around 60,000, a number that is, in all likelihood, a vast underestimate due to inadequate reporting requirements and *Leishmania* prevalence in areas with minimal access to healthcare (Alvar *et al*. 2012; Okwor and Uzonna 2016; Karimkhani *et al*. 2016; Bailey *et al*. 2017). Sandflies also transmit pathogens that cause severe diseases among which are three arboviruses (Sicilian virus, Naples virus, and Toscana virus; (Tesh *et al*. 1975; Batieha *et al*. 2000; Dionisio *et al*. 2003; Izri *et al*. 2008)), *Vesiculovirus* (Comer and Tesh 1991), *Orbivirus* (Depaquit *et al*. 2010; Phumee *et al*. 2021), *Flavivirus* (Acevedo and Arrivillaga 2008), and *Bartonella bacilliformis*, the etiological agent of Carrion’s disease (Caceres *et al*. 1997).

Species from the Psychodidae family occupy a variety of habitats. Psychodinae species are often associated with synanthropic habitats but are also commonly found in drains and aquatic environments (Cordeiro and Wagner 2018). Phlebotominae sandflies are often associated with places where they can obtain blood meals such as human settlements (Ramos *et al*. 2014) and coffee plantations (Ferro *et al*. 2015). Mating often occurs at lekking sites at the base of large trees (Memmott 1991, 1992). Females tend to oviposit close to these lekking sites in clay rich soils. Painstaking collections have revealed the presence of immature stages in such sites (Rutledge and Mosser 1972; Vivero *et al*. 2015) but the precise details of most species remain largely unknown (Alexander 2000). The specificity of habitat varies across species, and in some cases even populations, and while some species are geographically widespread and inhabit a variety of habitats others seem to be more restricted in their geographic and ecological distribution.

One of the potential ecological niches of sandflies is karstic landscapes (Oca-Aguilar *et al*. 2013; Blavier *et al*. 2019; Costa *et al*. 2021). Karstic landscapes are irregular limestone formations affected by erosion that encompass fissures, sinkholes, underground streams, and caverns. Previous reports have identified that some Phlebotomine species frequently breed in cavern systems. Quate (Quate 1962) carried out the first cave collection in the Batu Caves (Malaysia) and identified 22 phlebotomids from different subfamilies. Since then, it has become clear that cavern soil is a potential breeding ground for Psychodids, especially for Phlebotomines. In total, 161 species of Phlebotomines have been isolated from caves. Notably 37 Phlebotomine species were first described from collections in caves (e.g., (Quate 1962; Galati *et al*. 2003; Alves *et al*. 2011; Campos *et al*. 2017; Costa *et al*. 2021)) and several species are exclusively found in caves (Alves *et al*. 2008; Barata *et al*. 2012; Vilela *et al*. 2015). Breeding in caves is a trait that is geographically widespread in the family. Sixty-six of the recorded cave dwelling species are specific to the Old World, and 99 occur in the New world. Clearly, cave surveys have the potential to add crucial information about the diversity of sandflies.

South and Central America harbors a high diversity of Phlebotomine species (D’Agostino et al. 2022) and they house the vector genus *Lutzomyia sensu lato*. Samplings from Brazil suggest the existence of species restricted to caves (Alves *et al*. 2011; Carvalho *et al*. 2013; Campos *et al*. 2017), and that important disease vectors can breed in caves (Carvalho *et al*. 2017). Few efforts have addressed the diversity of this group in caves in the Neotropics, especially in Northern South America. The only study on this matter was incidental and found Phlebotomines associated with a karstic, not cavernous, system. Bejarano et al. (Bejarano *et al*. 2018) reported the presence of *Warileya (Hertigia) hertigi*, a Phlebotomine not involved in the transmission of disease. Addressing the diversity of Phlebotomines in caves in the Northern Andes holds the potential to reveal general patterns about the diversity of this vector group, which remains a largely unaddressed question.

We report the results of a point sampling of a cave system in the Los Guácharos Cave in the Andean forest at la Reserva Natural Cañón del Río Claro (Antioquia, Colombia). We report three species previously unknown to occur in Colombia, and four species that were not known to occur in karstic environments. We then used existing datasets to test the hypothesis that Neotropical cave sandfly species diversity follows the expectations of a latitudinal species richness gradient. We find that cave sandfly species diversity does not follow the expectations of a latitudinal species richness gradient and that the species diversity in tropics and subtropics seems to be similar. Finally, we used comparative phylogenetic methods to study whether association to karstic environments has evolved multiple times in the evolutionary history of the Psychodidae family. We find that association to karst is not a trait with a strong phylogenetic signal. Our findings suggest that the diversity of the family is largely understudied and will require the incorporation of cave systems.

## METHODS

### Locality

The focus of our collection was the cave system at the Cañón del Río Claro Nature Reserve. The reserve is located between the municipalities of Sonsón, Puerto Triunfo and San Luis, to the southwest of the department of Antioquia (5º 5 ‘N, 74º 39’ W, (Vivero *et al*. 2010)). This cave system is of karstic origin and was formed by the erosion of irregular limestone (Restrepo Martinez 2011). The largest cave of the cave system, the Los Guácharos cave, has a length of 442.8 meters, with an entrance 10 to 15 meters high by 2 to 3 meters wide and is considered a humid cave (Moncada et al., 1989). The environment inside the karstic formation can be classified as caverns, fissures, sinkholes, and underground streams. The cave serves as a roosting site for oilbirds (*Steatornis caripensis*) and bats.

### Sampling

To characterize the entomofauna in the cave, we carried out a sampling during two consecutive nights in the month of September 2019. We installed 10 CDC-type light traps for a period of 12 hours starting at 18:00. The traps were collected at 6:00 the following morning. To attract blood feeding insects, we placed dry ice under the traps. The sublimating CO_2_ mimics the respiration of vertebrates and increases the sandfly yield. Two of the CDC traps were located in a cave with a 2 m^2^ cavity. Two more were placed inside a karst fold with a depth of 4 meters and an altitude of approximately 4 meters. The other six traps were placed in the main gallery of the cavern. The location of the traps obeyed safety precautions and the depth of bodies of water (Figure 1). For comparison, we installed a CDC trap in the vegetation outside the cave. We compared the species richness outside and inside the cave.

**FIGURE 1.**
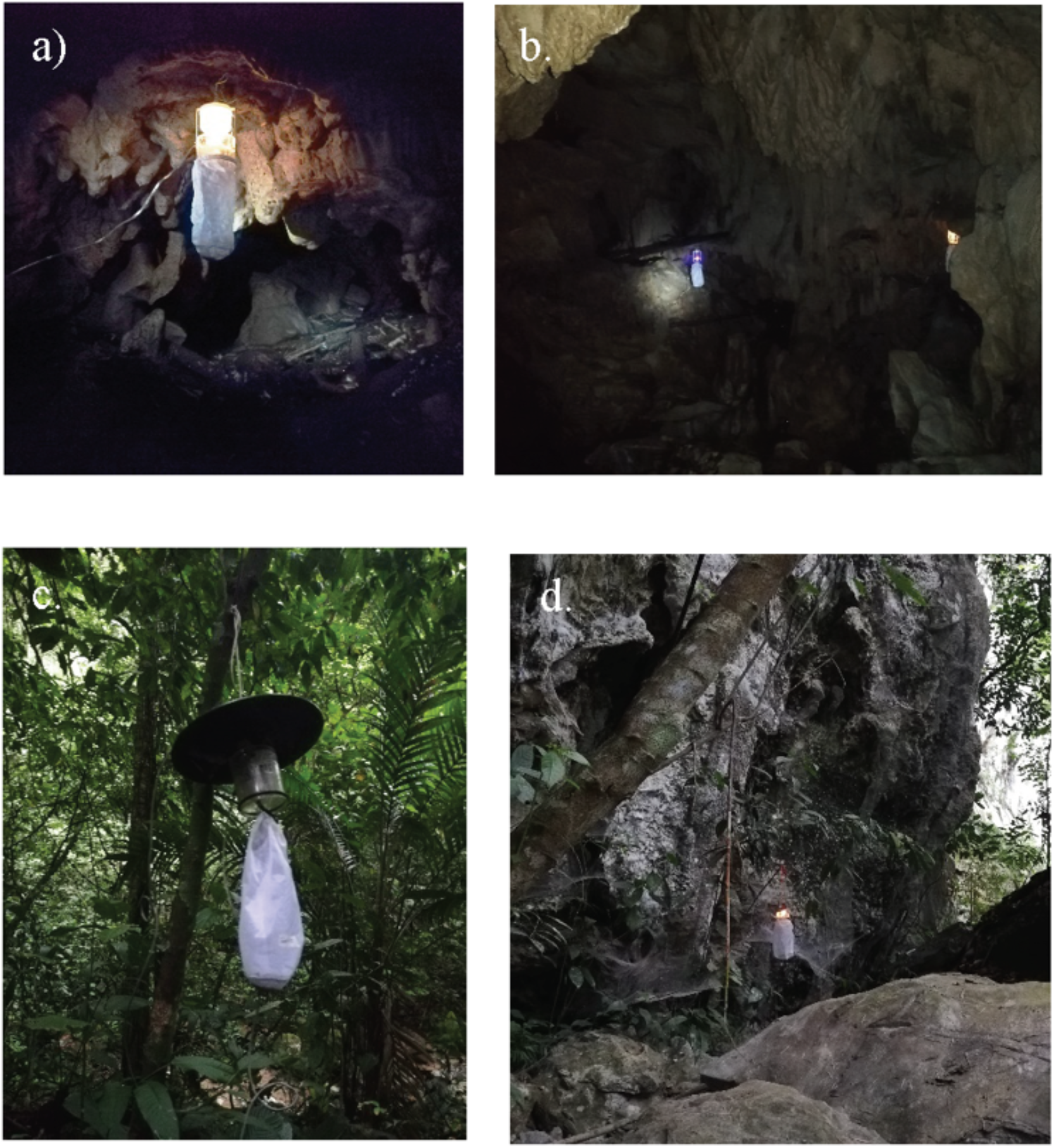
CDC light traps in the Los Guácharos cave collection. **A-B**. Cave. **C**. Vegetation outside the cave. **D**. Karstic folds.

For species identification, we transported the nets of the CDC traps to the Laboratory of Medical Entomology of the Program for the Study and Control of Tropical Diseases of the University of Antioquia. We extracted specimens from the traps using an aspirator and placed them in a Petri dish and inspected under a Leica microscope. To clear samples, we immersed each sandfly in lactophenol (1:1 ratio). Finally, we identified samples to species-level following the taxonomic classification scheme proposed by (Galati 2019).

### Species diversity and comparison to other studies

Next, we calculated metrics of species diversity in the Los Guácharos cave and compared them to other cave collections in the Neotropics to determine whether sandfly species richness followed a latitudinal gradient. Forty-six studies have reported the occurrence of Phlebotomines in caves. Thirty-six of them have addressed the taxonomic patterns of caves. We excluded samplings that just reported ecological diversity but did not include the raw data. After excluding these studies, we compiled data from 18 other cave samplings (listed in Table 1). Using this dataset, we calculated three different metrics of species diversity. First we used the number of collected species. Since the effort and size of the 19 samplings differed among studies, this metric is inherently limited. Second, we used the Shannon’s Index (*H*) which is a weighted geometric mean of species abundances. *H* follows the form:

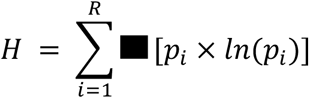

where *p*_*i*_ is the proportional representation of a given species and R is the number of species in the sample. If most of the abundance in the sample is concentrated in a single species, then H will be 0. In cases in which all the species are equally abundant, *H* will equal ln(R).

**TABLE 1.** Compilation of neotropical studies of sandfly cave diversity and their respective metrics of species diversity. H: Shannon’s H index. **\***Includes estimates over multiple years.

| Locality | Reference | H | Species richness | Evenness |
| --- | --- | --- | --- | --- |
| Los Guácharos Cave, Colombia | This study | 2.664 | 19 | 0.905 |
| Quadrilatero Ferreira, Brazil | (Campos *et al.* 2020) | 1.637 | 11 | 0.683 |
| Salitre, Brazil | (Campos *et al.* 2020) | 1.148 | 10 | 0.498 |
| Minas Gerais, | (Barata and | 1.432 | 20 | 0.478 |
| **Brazil** | Apolinário 2012) |  |  |  |
| **Bodoquera, Brazil** | (Galati *et al.* 2003) | 0.968 | 13 | 0.378 |
| **Pitangueiras, Brazil** | (Galati *et al.* 2003) | 0.171 | 12 | 0.069 |
| **Anhumas, Brazil** | (Galati *et al.* 2003) | 0.479 | 16 | 0.173 |
| **Guaicurus, Brazil** | (Galati *et al.* 2003) | 1.554 | 12 | 0.626 |
| **Jardim, Brazil** | (Galati *et al.* 2003) | 0.740 | 3 | 0.674 |
| **North Rondônia, Brazil** | (Ogawa *et al.* 2016) | 1.723 | 22 | 0.557 |
| **South Rondônia, Brazil** | (Ogawa *et al.* 2016) | 2.015 | 17 | 0.711 |
| **Gruta do Maruaga, Brazil** | (Alves *et al.* 2011) | 0.188 | 13 | 0.073 |
| **Gruta dos Lages, Brazil** | (Alves *et al.* 2011) | 0.436 | 8 | 0.210 |
| **Gruta dos Animais, Brazil** | (Alves *et al.* 2011) | 1.516 | 8 | 0.729 |
| **Millonario, Belize \*** | (Williams 1976) | 0.418 | 8 | 0.215 |
| **Augustine, Belize** | (Williams 1976) | 0.449 | 8 | 0.195 |
| **SA, Belize** | (Williams 1976) | 0.963 | 11 | 0.402 |
| **Yucatan, Mexico** | (Oca-Aguilar *et al.* 2013) | 1.180 | 7 | 0.607 |
| **Bambui, Brazil** | (Carvalho *et al.* 2013) | 1.903 | 22 | 0.616 |
| **Pains, Brazil** | (Campos *et al.* 2017) | 1.917 | 24 | 0.603 |
| **Pitoco, Brazil** | (de Almeida *et al.* 2019) | 1.927 | 22 | 0.623 |
| **Parauapebas, Brazil** | (Teodoro *et al.* 2021) | 1.353 | 17 | 0.477 |
| **Curionópolis, Brazil** | (Teodoro *et al.* 2021) | 1.771 | 13 | 0.691 |

We calculated a second index of diversity, evenness, which is defined as H normalized by the number of species and follows the form:

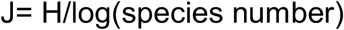

All metrics were calculated using the R package *vegan (*function *diversity*, (Oksanen 2013; Oksanen *et al*. 2013, 2015)).

We next studied whether there was a relationship between different metrics of cave sandfly species richness and latitude. Our expectation was that species richness in caves would follow a latitudinal pattern of diversity and that caves closer to the tropics would show higher species richness, as has been observed for other taxa (Mittelbach *et al*. 2007; Mannion *et al*. 2014; Jablonski *et al*. 2017; Pontarp *et al*. 2019). We regressed the different metrics of species richness with the latitude of the cave using the function *lm* (library *stats*; (R Core Team 2016)). We bootstrapped the regression coefficients using the function *Boot* (library *simpleboot*, (Peng 2008)).

### The evolution of cavernicolous niche

Our work and others report several unrelated species in the Pscychodidae family to have a cavernicolous niche. This suggests the possibility that the niche has evolved several times in the evolutionary history of the group. We formally addressed whether cavernicolous niche has evolved more than once in the evolutionary history of the family using comparative phylogenetic methods. We used a previously published phylogeny of the group using COI (D’Agostino et al. 2022) and classified each species as cavernicolous (i.e., has been collected in caves) or non-cavernicolous. We used the phylogeny and this classification to measure whether cave habitat followed a phylogenetic pattern. We used Pagel’s λ (Pagel 1999) for discrete traits (i.e., whether a species had been found in caves or not). Pagel’s *λ*, is a measure of phylogenetic signal which estimates the extent to which the phylogenetic history of a clade is predictive of the trait distribution at the tree tips. Values of λ lower than 1.0 represent traits being less similar amongst species than expected from their phylogenetic relationships. A λ equal to 1.0 suggests that traits covary with phylogeny (Pagel 1997, 1999) and is consistent with niche conservatism along the phylogeny (Cooper *et al*. 2010). A value close to zero indicates no effect of the phylogenetic history on the evolution of a trait. We used the wrapper *fitLambda* for the *ace* function (library *ape*, (Paradis *et al*. 2004; Paradis and Schliep 2019)) for ancestral character estimation and *λ* calculation. This function optimizes Pagel’s *λ* tree transformation for a discrete character evolving by a continuous-time Markov chain.We also used the function *fitdiscrete* (library *geiger*, (Harmon *et al*. 2008; Pennell *et al*. 2014)) which gave us the same results.

## RESULTS

### Collection description

We collected 52 sandflies, 16 males and 36 females. This difference in the sex ratio of the collection obeys different patterns of attraction to light traps (Toprak and Özer 2007). The samples belonged to 11 species of the Phlebotominae subfamily, representatives of the genera *Warileya, Lutzomyia, Micropygomyia, Bichromomyia, Evandromyia, Pintomyia, Pressatia, Pschoathyrdoomyia, Pschoathyromygusia*, and *Helcorcitomyia* (Table 2). The most abundant species was *Pintomyia ovallesi* (17.3%) followed by *Warileya hertigi* (11.53%) and *Lutzomyia gomezi* (11.53%; (Table 2)). The other species present in the cave environment correspond to *Warileya leponti* (9.61%), *Brumptomyia brumpti, Lutzomyia hartmanni* and *Psychodopygus panamensis*. This collection represents two novel findings. First, three species are novel for Colombia which significantly expands the range of these species. Second, we report four species that had not been, or have been rarely, collected in caves. We discuss these findings as follows.

**TABLE 2.**
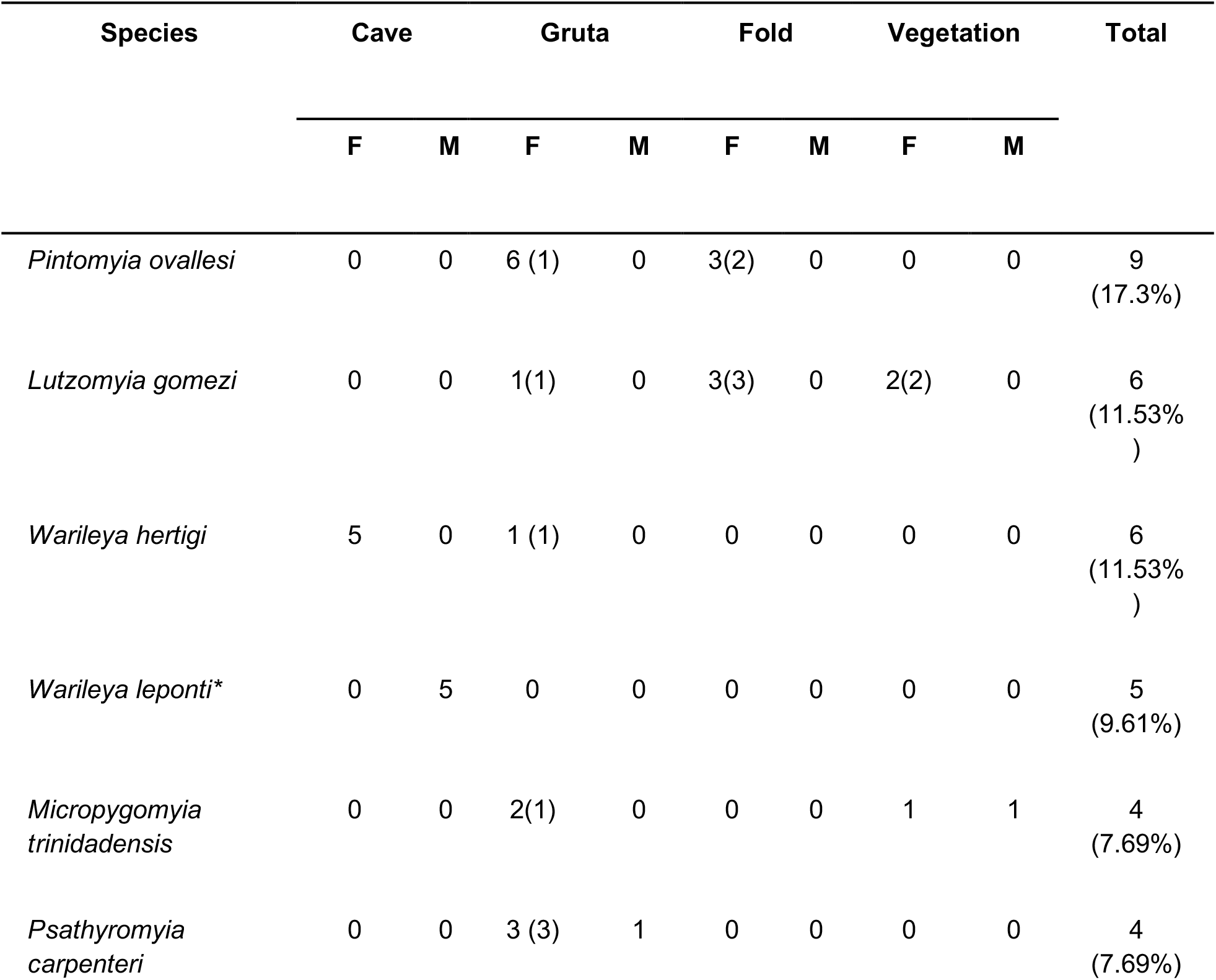
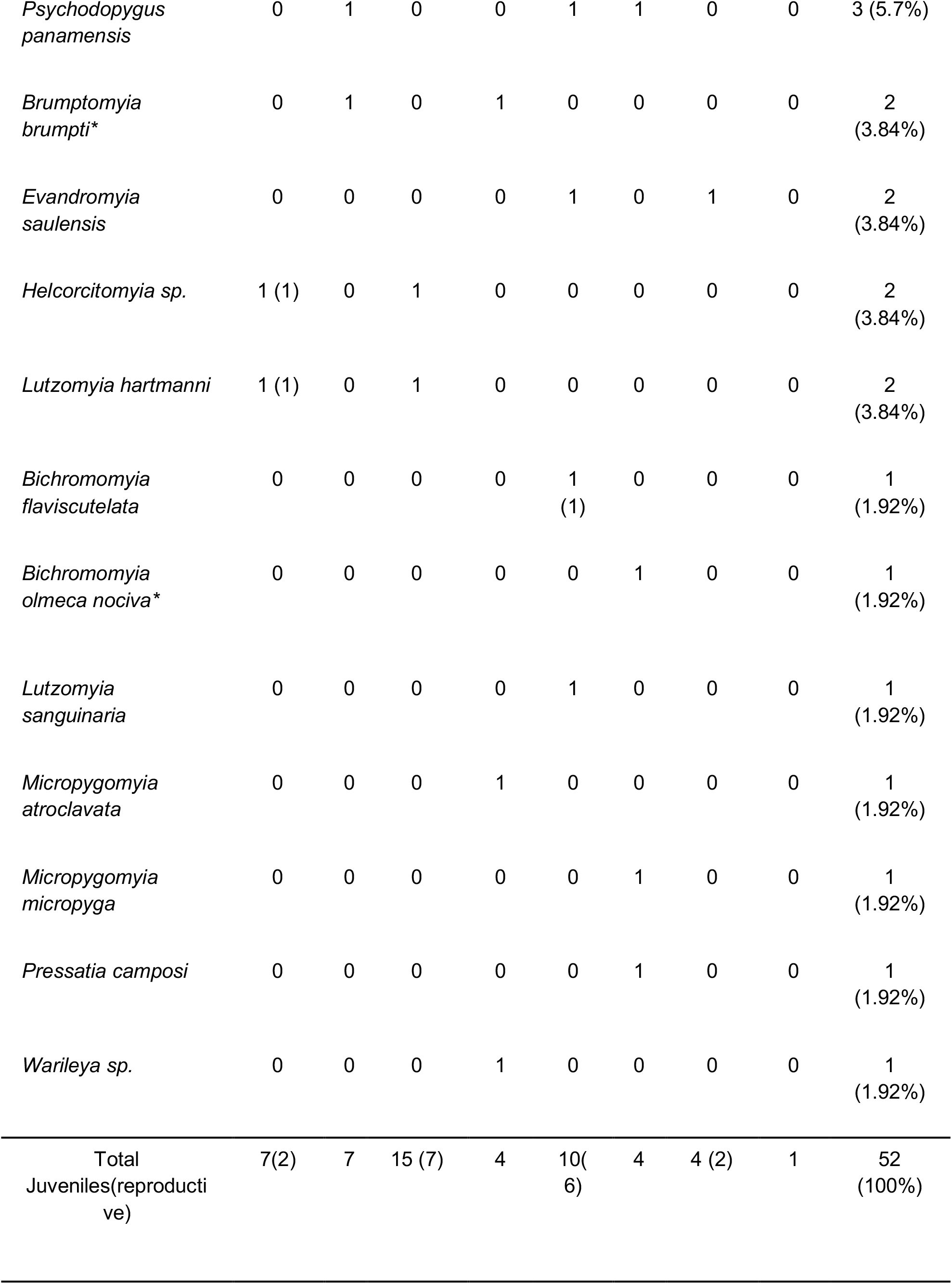
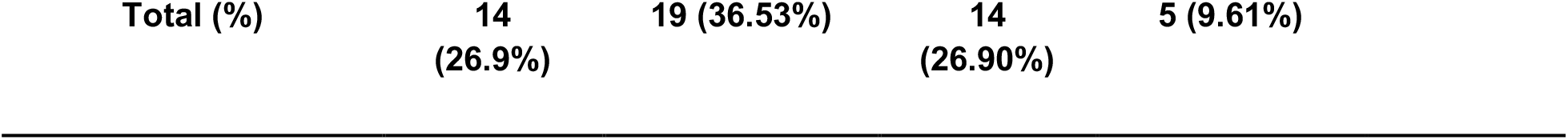
List of Phlebotomines collected in the Los Guácharos Cave in the Rio Claro Natural reserve (Antioquia), Colombia. F= female; M= male. * = species collected in Colombia for the first time. The number within the parentheses shows the number of gravid females or males with rotated genitalia (reproductive stage).

### New reports for Colombia

Our sampling includes three species that had not been reported for Colombia. We describe the morphological features of these specimens as follows.

#### Warileya leponti

(Galati & Cáceres, 1999): Cavern (5 ° 53’16.7384 “N, 74 ° 51’00.2101” W), Cañón del Río Claro Natural Reserve (Antioquia); Figure 2 A-C. (♂), fifth palpal segment is 0.8 mm; second palpal segment is 1.0 mm. Interocular suture (1 mm) is similar to the eye width (1 mm). Sperm pump (1.0 mm) is shorter than the aedegal ducts (2.0 mm), gonocoxite without tufted bristles and with approximately three isolated bristles. *Warileya leponti* has only been reported in Perú in the district of Villa Rica, province of Oxapampa (Pasco) (Cáceres *et al*. 2000); no systematic sampling exists for the species. To date, there were only records of *Warileya rotundipennis, Warileya nigrosacculus* and *Warileya hertigi* in Colombia (Bejarano 2006; Bejarano *et al*. 2018), which makes this collection of *W. leponti* the first for the country, and represents a large range extension for *W. leponti* (∼1,300 km North of previous collection).

**FIGURE 2.**
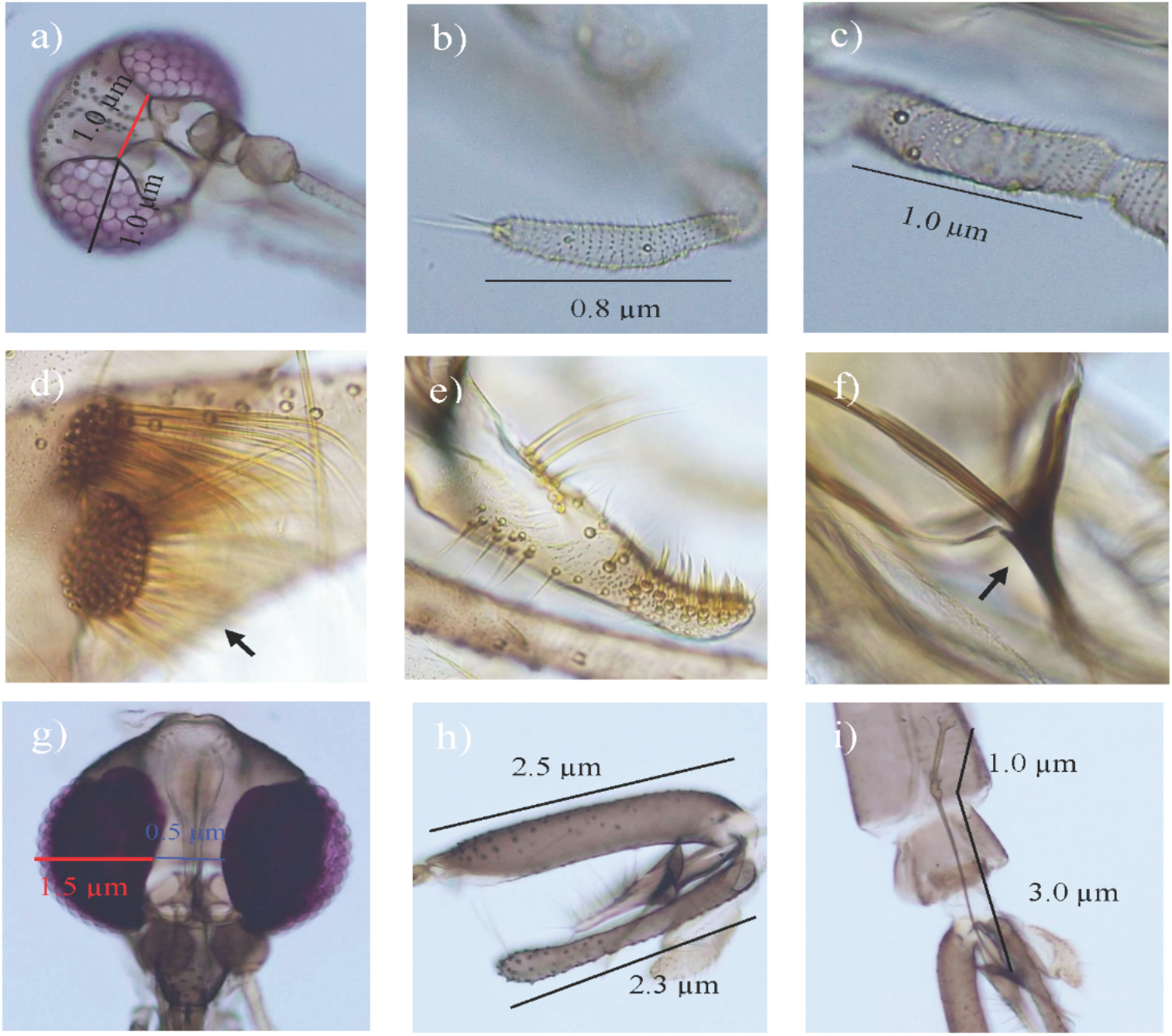
Diagnostic traits of the new Phlebotomine registers for the Colombian territory. *Warileya leponti*: (a) Head, complete interocular suture. Fifth (B) and second (c) palpal segments. *Brumptomyia brumpti*: (d), Tuft of simple setae on gonocoxite (e), paramere with inner spinelike-setae, (f) Aedeagus. *Bichromomyia olmeca nociva*: (g) Head and interocular space, (h) Epandrial lobes slightly shorter than gonocoxite, (i) spermiducts/sperm pump.

#### Brumptomyia brumpti

(Larrousse 1920): cavern (5 ° 53’16.8717 “N 74 ° 51’00.3429” W) and grotto (5 ° 53’17.6305 “N 74 ° 50’59.1009” W), Cañón del Río Claro Nature Reserve (Antioquia); Figure 2 D-F. (♂), gonocoxite with tuff formed by 20 semi-follicle bristles implanted directly on the surface, paramere without bifurcations with a dense group of fine and strong bristles located in their apical part and on the dorsal margin with 13 long bristles. Conical parameral sheath and gonocoxites without apical bristles and long basal bristles. *Brumptomyia brumpti* (Larrousse 1920) was found associated with the interior of the cave and in the closest cave located in the upper part of the cave. This species was previously reported only in Brazil (Galati, 2019), roughly 3,300 km southwest of our sampling.

#### Bichromomyia olmeca nociva

(Young and Arias 1982): folding (5° 53’ 16.8729 “N, 74 ° 51’ 17.6029” W), Cañón del Río Claro Nature Reserve (Antioquia); Figure 2 G-I. (♂), interocular distance (1.5 mm) equivalent to 1/3 of the width of the eye (0.5 mm), epandrial lobe (2.3 mm) shorter than the gonocoxite (2.5 mm), ratio of edegal ducts (3.0 mm) / sperm pump (1.3 mm) equals to 3.0:1.0. *Bichromomyia olmeca nociva* was found in the inner part of the cave across the river. Before this report, *B. olmeca nociva* has been reported for the Brazilian Amazon but not for the Colombian territory (∼1,700km extension).

### New reports for cave-dwelling species

Our sampling found four species that have either not been collected in caves, and one species which has only been reported in a cave once before. In this sampling, we found that *Lutzomyia hartmanni* (Fairchild and Hertig 1957) appeared associated with two different karstic environments, caves and rock-breaks. *Lutzomyia gomezi* (Nitzulescu 1931) has not been reported before in association with caves or any karstic systems. However, in this report we find the species in the Los Guácharos cave and surrounding vegetation. In both instances we found females that recently had a blood meal. Finally we observed two more species, *Psychodopygus panamensis* (Shannon 1926) and *Pintomyia ovallesi* (Ortiz 1951) in caves for the first time in our sampling.

*Psychodopygus panamensis* is a generalist species that is often characterized as anthropophilic (González *et al*. 1999; Santamaría *et al*. 2020; Rigg *et al*. 2021) as is *Pintomyia ovallesi* (Feliciangeli 1997; Feliciangeliand Rabinovich 1998; Rabinovich and Feliciangeli 2004). One previous study has collected samples from this species in a cave in Belize (Williams 1976). In both the Belize collection and our collection, females of *Psychodopygus panamensis* had blood-fed. Finally, we found *Bichromomyia flaviscutelata* (Mangabeira Filho 1942) in the Los Guácharos cave, agreeing with a previous report in Brazilian caves that reported this species can have a cavernicolous habitat (Ribeiro 2013).

### Species richness and comparisons to other caves

Our sampling yielded 52 individuals and revealed the existence of 18 species in the Los Guácharos cave. This sample has the highest species richness in any Neotropical sample collected in caves (Table 1). Nonetheless, our sample has a relatively low sample size. To account for differences in the sample size we calculate a Shannon diversity index and a normalized evenness index for each of the 19 caves sampled in the neotropics. The Guacharos cave also has the highest H and evenness of any neotropical cave suggesting the highest diversity of sandflies in Neotropics to date. Regardless of the species metric diversity, the Los Guácharos cave has high species diversity compared to other samples.

Next, we tested whether the diversity of sandflies in caves followed a latitudinal gradient of diversity by regressing the latitude of the collection with each of the three metrics of diversity (Figure 3). Contrary to our expectation, we find that none of the three regressions is significant suggesting that species diversity of sandfly caves does not decrease as the cave gets farther from the Equator (F_1,21_ < 0.229, P > 0.637).

**FIGURE 3.**
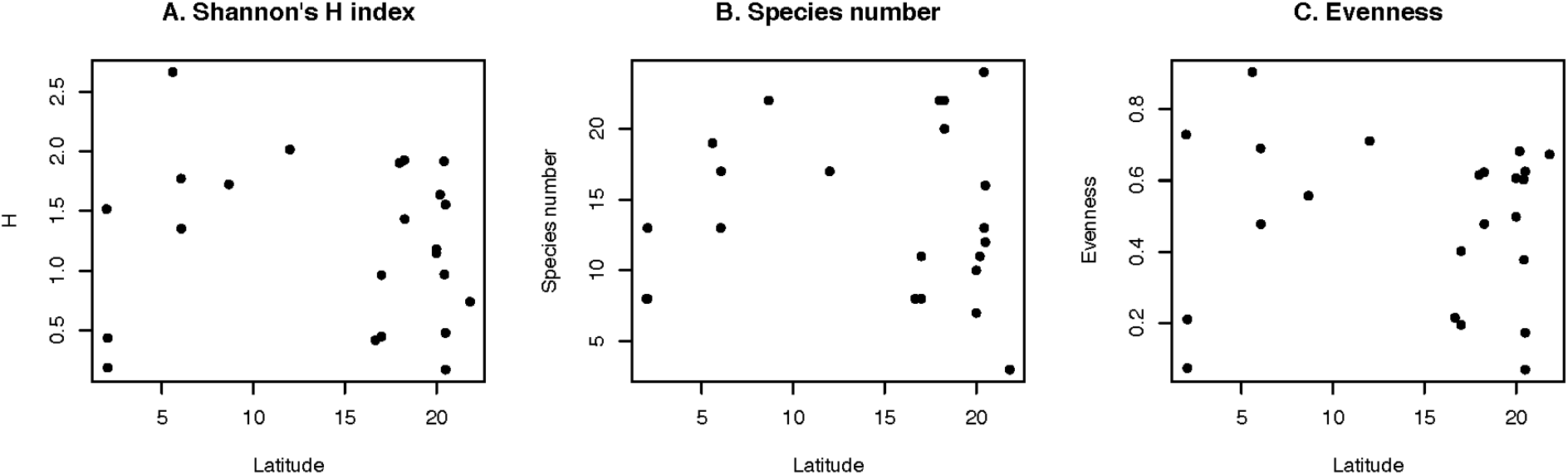
Three different metrics of the species diversity in Neotropical sandfly cave collections. **A**. Number of species. **B**. Shannon’s *H* index. **C**. Evenness. Each dot represents the species richness metric for a sampling listed in Table 1. None of the three metrics shows evidence for a latitudinal gradient of sandfly species diversity collected in caves.

### Phylogenetic signal of cavernicolous habitat

We used a COI genealogy to study whether the cavern habitat association had a phylogenetic signal in the family. Our metric of phylogenetic signal was close to zero (Pagel’s λ = 5.729 × 10^-5^; logLik = -42.95) which suggests no effect of the phylogeny on the distribution of cavernicolous habitat. The distribution of the trait along the tree suggests the same pattern as cavernicolous species are not related to each other (Figure 4). Thus cave use likely evolved at least three times in the family as it appears in at least three different genera.

**FIGURE 4.**
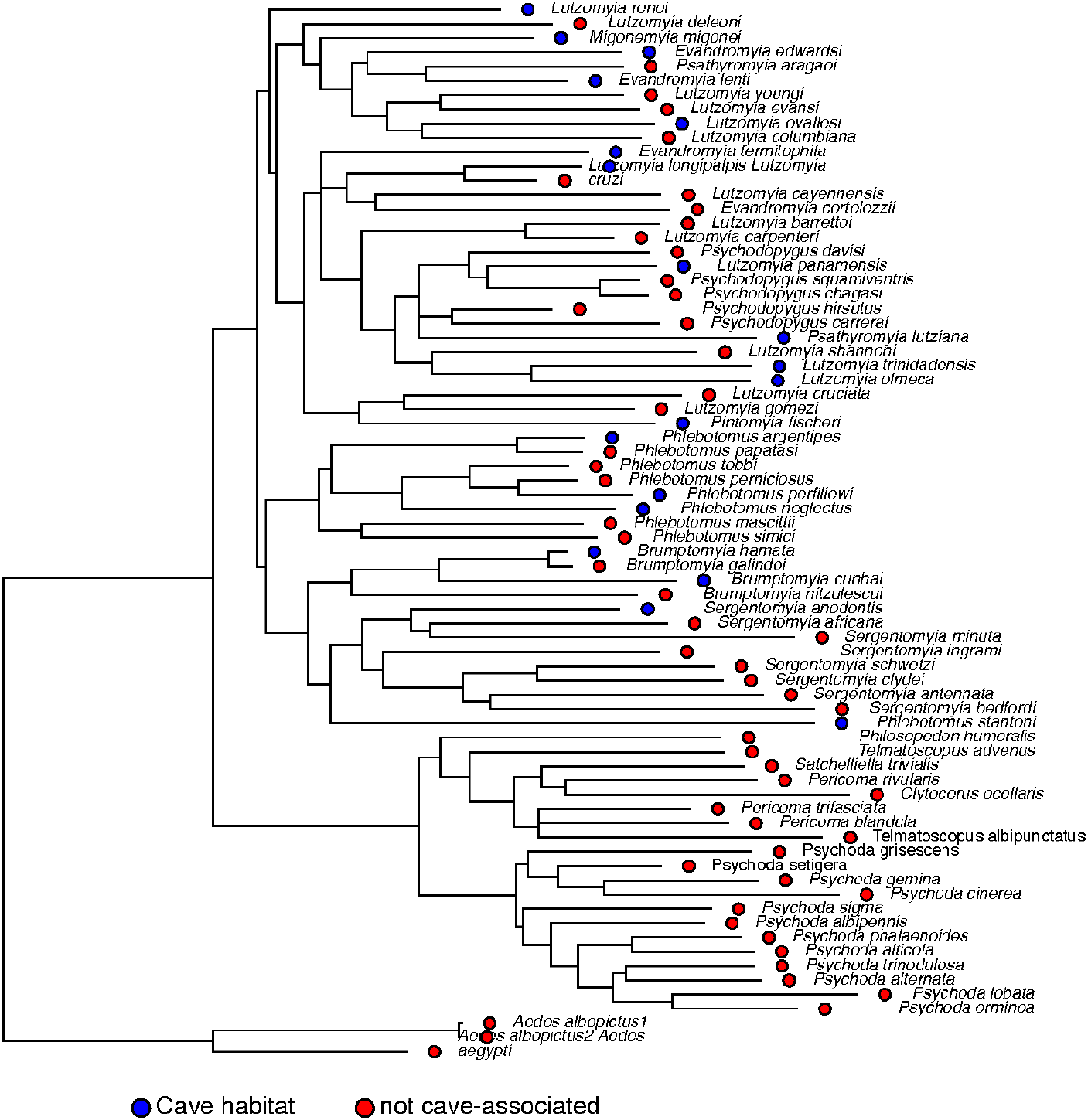
Phylogenetic signal of cavernicolous habitats in Psychodidae. The dots at the tree tips show the extant known habitat choice for each species in the phylogeny. Blue: known to be associated with caves, Red: not associated with caves.

## DISCUSSION

The study of the habitat of vector species can yield important information on potential control strategies to limit their ability to transmit disease. Nonetheless, the habitat of many vector species remains largely unknown. Among vectors, this knowledge gap is particularly problematic for sandflies of the phlebotominae tribe. This taxa includes vector species of human pathogens such as *Leishmania* (*Lutzomyia, Phlebotomus*) and species known to harbor pathogens but not transmit disease to humans (*Sergentomya, Warileya, Brumptomyia*). Species in the tribe occupy a wide variety of habitat ranges but the precise habitat of most species remains unknown (e.g., (Souza *et al*. 2017; Sandoval-Ramírez *et al*. 2020)). Moreover, some species have been reported in habitats where accessibility and sampling is difficult, such as caves. We did a point sampling of a cave in western Colombia. Our report is the first systematic *Lutzomyia* sampling of a cave in the northern Andes.

Our sampling revealed the presence of species in Colombia that were not known to occur in the country. *Warileya leponti* and *Brumptomyia brumpti* are two species that can harbor *Leishmania* but do not transmit it to humans. The biology of *Warileya* is largely unknown. The genus has been associated with environments with little or no human intervention, highlighting its importance as bioindicators of ecosystems (Vivero *et al*. 2017). Although the species of this genus are not related to the transmission of *Leishmania*, recently, *Warileya rotundipennis* was found with natural infection of parasites similar to *Leishmania* (*Viannia*) in Colombia (Moreno *et al*. 2015). Whether *W. leponti* shows natural infections remains unexplored.

*Brumptomyia brumpti* has collection records in Ecuador and Brazil (GBIF: DOI10.15468/dl.fhtqq8, years: 1959-2013). Previous reports have found the species associated with cave environments, at the entrance and inside the caverns (Campos *et al*. 2017), consistent with the finding of this study. *Brumptomyia brumpti* is a synanthropic species due to its high adaptive capacity, being found in both forested and disturbed environments (Capucci 2021).

A third species, *Bichromomyia olmeca*, has vector importance. The species is naturally infected and can effectively transmit *Leishmania amazonensis* (Barroso, 2019), which highlights the importance of finding this species in cave systems relatively close to human settlements and with frequent ecotourism. *Bichromomyia olmeca* seems to be a species complex composed of at least four different lineages: *B. o. nociva* (reported in this study), and *B. o. olmeca* (mainly localized in Mexico), *B. o. bicolor* (reported in Pánama, Colombia, and Brazil), and *B. flaviscutellata sensu stricto* (de Melo *et al*. 2020). The species complex is found throughout Central America to Brazil (Galati, 2019) (GBIF: DOI: https://doi.org/10.15468/dl.g77tbs, years: 1953–2018 and Table S1 in (Moo-Llanes *et al*. 2013)) and in some instances–as is the case in the Los Guácharos cave–, lineages have been found to coexist in the same geographic range (Fairchild and Theodor 1971; Escobar Vasco 1989; Romero Ricardo *et al*. 2013; de Melo *et al*. 2020; Santatana *et al*. 2020). The identification of Colombian isolates of this species poses the need to revisit the geographic range of each species, with a special emphasis on areas of coexistence which might facilitate gene flow between vector species.

Determining range limits has practical implications for our understanding of vector biology. Under current global warming scenarios, *B. olmeca olmeca* is predicted to expand its current range to subtropical and temperate areas (Moo-Llanes *et al*. 2013). This prediction was only possible after a systematic habitat characterization of the species habitat (108 collection sites, Table 1 in (Moo-Llanes *et al*. 2013)) which in turn revealed the suitable current and future ecological niche of the species. To date the habitat characterization of *B. olmeca nociva* or other species in the species complex, at least to the same detail as *B. o. olmeca*, is lacking. We hypothesize that lack of systematic sampling explains the discontinuous geographic range of other species (e.g., the substantial range extensions of *Bichromomyia olmeca nociva, Brumptomyia brumpti*, and *W. leponti* in this study).

More generally, we used the data from our sampling to infer ecological and evolutionary patterns about cave habitat use in sandflies. We find that the Los Guácharos cave has the largest species diversity of any sampled Neotropical cave. Opposite to our expectations, this analysis revealed that sandfly cave species diversity does not follow a latitudinal gradient in the South American continent. Latitudinal gradients, in which the tropics are the most diverse locales, have been reported for almost all systematically sampled taxa (reviewed in (Mannion *et al*. 2014; Jablonski *et al*. 2017); exceptions listed in (Kindlmann *et al*. 2007)). Nonetheless, the drivers of the pattern are hotly debated and are likely to be caused by a variety of processes. Fish for example show higher diversity in the tropics (Mittelbach *et al*. 2007; Stuart-Smith *et al*. 2013) but a higher speciation rate in temperate areas (Rabosky *et al*. 2018). Our results find that the variance of species richness might be the largest in the tropics, as the samples close to the equator show the highest (Los Guácharos cave, Colombia; this study) and some of the lowest species richness (Gruta dos Animais, Amazonas, Brazil, (Alves *et al*. 2011)) of all the sampled caves. More systematic sampling will be needed before the hypothesis that the tropics have a larger variance in species richness across localities but the possibility is tantalizing.

Finally, we assessed the evolution of cave habitat choice. We formally tested whether the occupation of cave habitats has evolved multiple times or just once in the Psychodidadae family. We find no phylogenetic signal for cave-habitat association which supports the possibility that habitat association has evolved repeatedly in the family. A different possibility is that many species are generalists that can use caves as a habitat when present, but limited karstic availability and/or sampling make verifying this difficult. In this scenario, species are only cavernicolous when there are available caves nearby, and the lack of phylogenetic signal in the trait suggests no genetic component to the trait. The current sampling does not allow us to differentiate between these possibilities. Mark recapture experiments releasing sandflies at the entrance of caves and determining their habitat choice should reveal whether the collection of species in caves is due to happenstance. One important caveat is that we expect the phylogeny of sandflies to change as data for more species and better genetic markers become available. The current topology is based on a single mitochondrial marker (COI) which has important limitations (reviewed in (Rubinoff and Holland 2005)). Therefore, we expect our estimates of phylogenetic signal to change slightly. However, our conclusion that cavernicolous habitat has evolved in several instances in the phylogeny will stand unless the topology is dramatically revised.

Few studies have done the painstaking collectections needed to address the specific habitats of immature stages of sandflies. Vivero et al. (Vivero *et al*. 2015) extracted 160 soil samples from a variety of habitats, including urban and undisturbed environments, and found sandflies in a surprising 35% of the samples. The study found at least 26 species of trees (Table 3 in (Vivero *et al*. 2015)) serve as breeding ground for *Lutzomyia* and two for *Brumptomyia*. A subsequent analysis (Estrada *et al*. 2020), focused on a single species, *Lu. evansi*, to explore urban and rural microhabitats and the association between this species and different tree substrates. While *Lu. evansi* is associated with multiple tree species, it is often associated with the base of mature *Cordia* trees. These assessments exemplify the difficulty of identifying the habitat of vectors and show why the topic remains largely understudied. Understanding the precise habitat tolerance and choice represents a crucial step to understand why some species are vectors while others are not.

We examined the sandfly diversity of a Northern Andean cave for the first time, and recorded three new records for Colombia, and four species that have previously not been associated with caves. Further, we examined the evolution of cave use within Psychodidae and discovered that the use of caves likely evolved repeatedly. Our results confirm the need for thorough and systematic sampling of sandfly diversity, including habitats that have not often been sampled (e.g., caves), in order to better understand this ecologically and medically important clade.

## ACKNOWLEDGEMENTS

We thank the Matute lab for constructive feedback on the manuscript. This work was supported by the National Science Foundation (Dimensions of Biodiversity award 1737752 to D.R.M.). The funders had no role in any aspect of study design, data collection and analysis, or decisions with respect to publication.

